# Improved Prokaryotic Gene Prediction Yields Insights into Transcription and Translation Mechanisms on Whole Genome Scale

**DOI:** 10.1101/193490

**Authors:** Alexandre Lomsadze, Karl Gemayel, Shiyuyun Tang, Mark Borodovsky

**Affiliations:** Wallace H. Coulter Department of Biomedical Engineering; School of Computational Science and Engineering; School of Biological Sciences, Georgia Tech, Atlanta, Georgia, 30332, USA; Department of Biological and Medical Physics, Moscow Institute of Physics and Technology, Moscow, Russia

## Abstract

In a conventional view of the prokaryotic genome organization promoters precede operons and RBS sites with Shine-Dalgarno consensus precede genes. However, recent experimental research suggesting a more diverse view motivated us to develop an algorithm with improved gene-finding accuracy. We describe GeneMarkS-2, an *ab initio* algorithm that uses a model derived by self-training for finding species-specific (native) genes, along with an array of pre-computed *heuristic* models designed to identify harder-to-detect genes (likely horizontally transferred). Importantly, we designed GeneMarkS-2 to identify several types of distinct sequence patterns (signals) involved in gene expression control, among them the patterns characteristic for leaderless transcription as well as non-canonical RBS patterns. To assess the accuracy of GeneMarkS-2 we used genes validated by COG annotation, proteomics experiments, and N-terminal protein sequencing. We observed that GeneMarkS-2 performed better on average in all accuracy measures when compared with the current state-of-the-art gene prediction tools. Furthermore, the screening of ∼5,000 representative prokaryotic genomes made by GeneMarkS-2 predicted frequent leaderless transcription in both archaea and bacteria. We also observed that the RBS sites in some species with leadered transcription did not necessarily exhibit the Shine-Dalgarno consensus. The modeling of different types of sequence motifs regulating gene expression prompted a division of prokaryotic genomes into five categories with distinct sequence patterns around the gene starts.

[Supplemental material is available for this article].

## Introduction

Since the number of microbial species on Earth is estimated to be larger than 10^12^ (Locey and Lennon 2016), the exponential growth of the number of sequenced prokaryotic genomes, currently ∼10^5^, is likely to continue for quite a while. Along this path, we will continue to see genomes with large numbers of genes not detectable by mapping protein orthologues. Thus, improving the accuracy of *ab initio* gene prediction remains an important task.

The proliferation of RNA-Seq presented an opportunity for more accurate inference of exon-intron structures of eukaryotic genes. Transcriptomes of prokaryotes, however, were thought to be less important for gene finding since the accuracy of *ab initio* prediction of a whole gene (being uninterrupted ORF) is significantly higher. Nevertheless, recent innovations in the NGS techniques led to generating new kinds of data whose impact has yet to be fully appreciated.

For example, dRNA-Seq, the differential RNA sequencing technique (Sharma and Vogel 2014; Sharma et al. 2010) aimed to accurately identify transcription start sites (TSS). The experimental evidence for the TSS locations is crucial for reliable operon annotation, as well as the detection of promoters and translation initiation sites (TIS) (Creecy and Conway 2015).

The sequence downstream from TIS is supposed to code for interactions between mRNA and the translation machinery. In prokaryotes, translation initiation is assumed to be facilitated by the base-pairing between the 3’ tail of the 16S rRNA of the 30S ribosomal subunit and the ribosome binding site (RBS) located in the 5’ UTR of the mRNA. The pioneer work of Shine and Dalgarno on *Escherichia coli* described a specific RBS consensus observed later in many prokaryotic genomes (Shine and Dalgarno 1974; Barrick et al. 1994). Still, with the accumulation of genomic data, exceptions started to multiply. For instance, the 5’ UTR may be completely absent in the case of *leaderless transcription*, first discovered in the archaea *Pyrobaculum aerophilum* (Slupska et al. 2001).

Recent studies of prokaryotic transcriptomes, including dRNA-Seq applications, detected instances of leaderless transcription not only in archaea but in bacteria as well (Cortes et al. 2013). Importantly, the fraction of genes with leaderless transcription was observed to vary significantly among species. It was low (<8% among all operons) in some bacteria, such as *Helicobacter pylori* (Sharma et al. 2010), *Bacillus subtilis* (Nicolas et al. 2012), *Salmonella enterica* (Kroger et al. 2013), *Bacillus licheniformis* (Wiegand et al. 2013), *Campylobacter jejuni* (Dugar et al. 2013), *Propionibacterium acnes* (Pfeifer-Sancar et al. 2013), *Shewanella oneidensis* (Shao et al. 2014), and *Escherichia coli* (Thomason et al. 2015). It was also low (<15%) in some archaea e.g. in *Methanosarcina mazei* (Jager et al. 2009), *Pyrococcus abyssi* (Toffano-Nioche et al. 2013), *Thermococcus kodakarensis* (Jager et al. 2014), *Methanolobus psychrophilus* (Li et al. 2015), and *Thermococcus onnurineus* (Cho et al. 2017). However, a higher frequency (>25%) of leaderless transcription was observed in other bacteria, e.g. *Mycobacterium tuberculosis* (Cortes et al. 2013), *Corynebacterium glutamicum* (Pfeifer-Sancar et al. 2013), *Deinococcus deserti* (de Groot et al. 2014), *Streptomyces coelicolor* (Romero et al. 2014), *Mycobacterium smegmatis* (Shell et al. 2015), and an even larger frequency (>60%) was seen in various archaeal species e.g. *Halobacterium salinarum* (Koide et al. 2009), *Sulfolobus solfataricus* (Wurtzel et al. 2010), and *Haloferax volcanii* (Babski et al. 2016). Accordingly, the diversity of regulatory sequence patterns that appear near gene starts motivated the effort to build multiple models necessary for more accurate gene-start prediction.

Current prokaryotic gene finding tools, GeneMarkS, Glimmer3, and Prodigal are known for a sufficiently high accuracy in predicting protein-coding ORFs. Indeed, on average these tools are able to find more than 97% of genes in a verified test set in terms of correct prediction of the gene 3’ ends (Besemer, Lomsadze, and Borodovsky 2001; Delcher et al. 2007; Hyatt et al. 2010). Furthermore, the accuracy of pinpointing gene starts is on average ∼90% (Hyatt et al. 2010). We observed that most of the genes that escaped detection altogether (false negatives) belonged primarily to the atypical category, i.e. genes with sequence patterns not matching the species-specific model trained on the bulk of the genome (Borodovsky et al. 1995).

Given the high accuracy of the current tools, the task to improve prokaryotic gene finding is challenging. That said, we describe below a method that not only improves the accuracy of gene prediction across the wide range of prokaryotic genomes, but also identifies genome-wide features of transcription and translation mechanisms.

## Methods

### Gene and Genome Modeling

#### Model of a protein-coding sequence

GeneMarkS-2 uses a rather complex model of a gene, a building block of the model of a prokaryotic genome (Fig. 1). The majority of protein-coding regions in prokaryotic genomes are known to carry species-specific oligonucleotide (e.g. codon) usage patterns (Fickett and Tung 1992). To that effect, GeneMarkS-2 learns this pattern and estimates the parameters of the *typical* model of protein-coding regions, a three-periodic Markov chain (Borodovsky et al, 1986), by iterative self-training on the whole genome, a procedure similar in general to the one introduced in GeneMarkS (Besemer, Lomsadze, and Borodovsky 2001).

**Figure 1.**
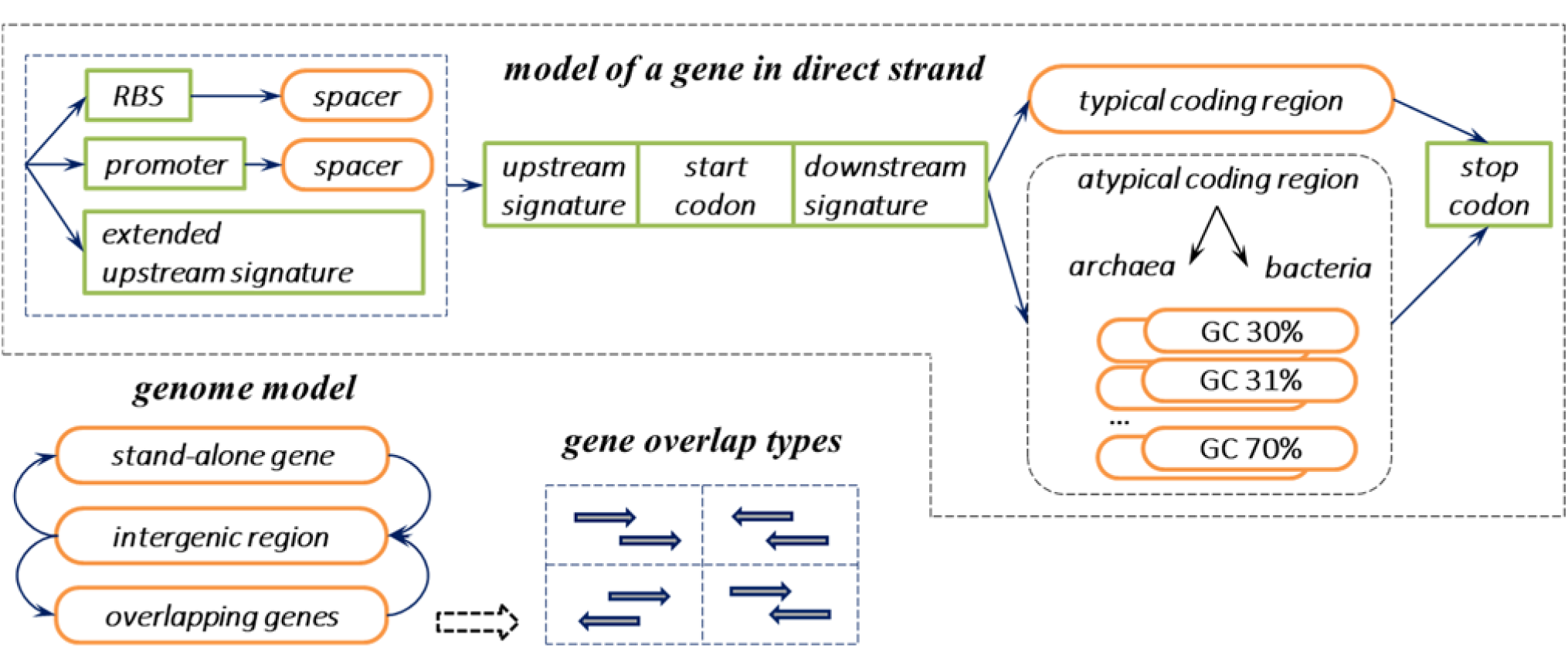
Principal state diagram of the generalized hidden Markov model (GHHM) of prokaryotic genomic sequence. States shown in the top panel were used to model a gene in the direct strand. Genes in the reverse strand were modeled by the identical set of states (with directions of transition reversed). The states modeling genes in direct and reverse strands were connected through the intergenic region state as well as the states of genes overlapping in opposite strands.

Still, the oligonucleotide composition of some genes may deviate from the genome-wide mainstream. For that reason, GeneMarkS introduced an additional *atypical* gene model alongside the *typical* model (Besemer, Lomsadze, and Borodovsky 2001). Instead of a single atypical model, GeneMarkS-2 uses two large sets of atypical models: 41 bacterial and 41 archaeal. The precomputed parameters of the atypical models, covering the GC content range from 30% to 70%, were estimated by a method utilized earlier in the metagenome gene finder (Zhu, Lomsadze, and Borodovsky 2010); these parameters are not re-estimated in GeneMarkS-2 iterations. Each atypical model has a GC content index indicating a narrow (1%) GC bin that it represents. The GC of the candidate ORF is used to identify which of the atypical models should be deployed for that ORF analysis. As such, only a subset of the atypical models is used in a GeneMarkS-2 run on any given genome.

This multi-model approach could be interpreted in the following way. Disregarding for a moment the linear connectivity of genes in a given genome, we can think of this set of ‘disjoint’ genes as an instance of a small ‘metagenome’. The approach developed earlier for metagenome analysis (Zhu, Lomsadze, and Borodovsky 2010) employed a variety of models for the analysis of sequence fragments with varying GC contents; we use this library as the set of *atypical* models for GeneMarkS-2. Furthermore, the typical genes, making the majority of this disjoint gene set, could be effectively clustered and processed together to determine parameters for the *typical* model. With this set of models at hand, a given ORF is predicted as a gene by the best fitting model, i.e. the model (the *typical* or the GC-matching *atypical*) that yields the highest score.

#### Model of a sequence around the gene start

Sequence patterns near gene starts are species-specific; nevertheless, we identified groups of genomes with similar patterns that arguably were selected in evolution due to the common features of translation and transcription mechanisms. We have introduced four distinct categories/groups of the pattern models (A through D). We also observed that some genomes did not fall into one of these four groups; as such, we created the fifth category/group X to which all such genomes were assigned. In general, category X either indicates very weak regulatory signals (hard to detect and classify) or possibly signals related to a new and yet poorly characterized translation initiation mechanism.

The first, group A, represents genomes with an observed dominance of RBS sites having the Shine-Dalgarno (SD) consensus; the usage of leaderless transcription was negligible or nonexistent in such genomes. Genomes of the second category, group B, carry RBS sites with a “non-Shine-Dalgarno” (non-SD) consensus. Next, there are group C and group D which represent bacterial and archaeal genomes, respectively, characterized by a significant presence of leaderless transcription. Importantly, in group C genomes, the bacterial promoter signal (Pribnow box) is located at ∼ 10 nt distance from the start of the first gene in operon (transcribed in the leaderless fashion). In group D, the archaeal promoter box is situated at ∼ 26 nt distance from the starts of genes with leaderless transcription. Furthermore, other genes (i.e. internal genes in operons or first-genes-in-operons with *leadered* transcription) in genomes of groups C and D are likely to use RBS sites; we therefore use *dual gene start models* in these types of genomes. In group X, we saw weak, hard-to-classify regulatory signal patterns; still, some genes in genomes of group X could have an RBS signal with SD consensus.

The sites of promoter boxes as well as the non-Shine-Dalgarno (non-SD) and the Shine-Dalgarno (SD) RBS sites are separated from TIS by sequences with variable length (spacers). Therefore, the model of a regulatory signal should include the model of the site as well as the length distribution of the spacer (Fig. 1). We also explicitly modeled the triplet upstream to the start codon as a 3 nt long *upstream signature*. In addition, we also used a *downstream signature model* (Fig. 1) that captured the patterns within the short sequence (with length up to 12 nt) located immediately downstream from the start codon (Shmatkov et al. 1999).

For genomes in group X, we used a combination of the site model for RBS (if a very weak but nevertheless statistically significant RBS model can be found) and a 20 nt long *extended upstream signature* (Fig. 1) in an attempt to capture some patterns not represented by the models developed for groups A through D.

A brief outline of groups A-D, X is given in Table 1.

**Table 1.** Features of the regulatory site models used for genomes of groups A-D and X. A dash indicates that a particular module was not used; -26 and -10 indicate average nt distance between the promoter Pribnow box and the position of translation start.

| Groups | Transcription Features & Domain | Representative Species | RBS Consensus | Promoter Box Localization | Extended Upstream Signature |
| --- | --- | --- | --- | --- | --- |
| A | Leadered with predominantly SD RBS | *Escherichia coli* | SD | -- | -- |
| B | Leadered with predominantly non-SD RBS | *Bacteroides ovatus* | non-SD | -- | -- |
| C | Leaderless & Bacteria | *Mycobacterium tuberculosis* | SD | --10 | -- |
| D | Leaderless & Archaea | *Holobacterium salinarum* | SD | --26 | -- |
| X | Unclassified | *Synechocystis* | SD | -- | 20nt |

### Unsupervised training

The unsupervised training algorithm makes several two-step iterations (Fig. 2). Each iteration results in i/ genome segmentation into protein-coding (CDS) and non-coding regions (gene prediction) and ii/ model parameters re-estimation (Besemer, Lomsadze, and Borodovsky 2001).

**Figure 2.**
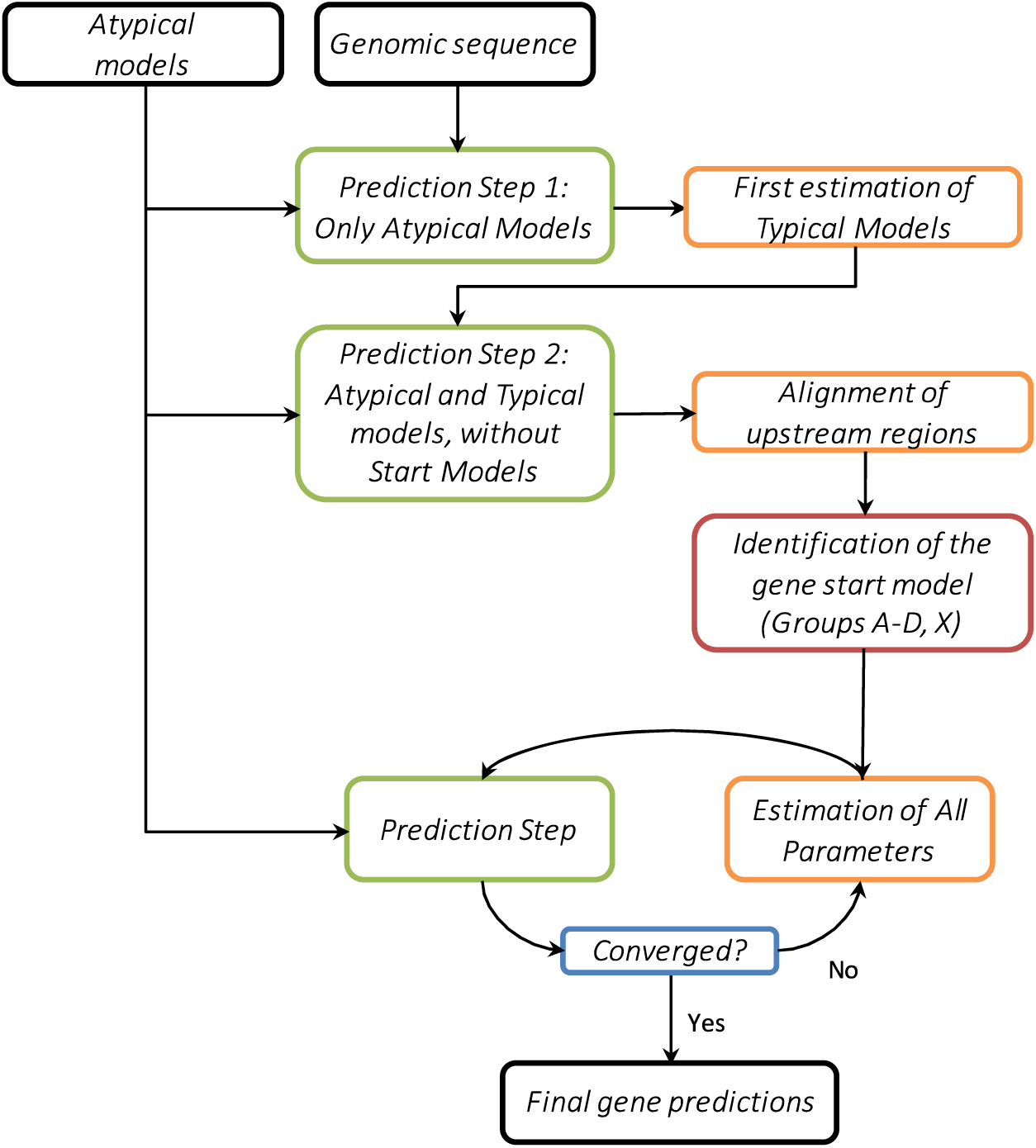
Principal workflow of the unsupervised training.

#### The first iteration

##### Prediction step

The log-odds space Viterbi algorithm (see Suppl. Materials) computes the maximum likely sequence of hidden states (Fig. 1) along the genome, the sequence delivering the largest log-odds score. At the first step, the algorithm only uses heuristic (atypical) models. When the algorithm runs on a particular segment of the genomic sequence, it utilizes only two of the 82 possible *atypical* models; specifically, the two, bacterial and archaeal, whose GC index matches the local sequence’s GC composition. In the first iteration, the algorithm does not account for sequence patterns near gene starts since no gene-start model has been derived yet.

##### Estimation step

After the first run of the Viterbi algorithm, all genomic segments predicted and labeled as ‘protein-coding’ are collected into a training set to estimate the parameters of the ‘typical’ model. To reduce the chance of including non-coding ORFs into the training set, predicted genes shorter than 300 nt and those predicted as incomplete CDS are not used in training. From thus complied sequence set, the *typical* model, a 5^th^ order three-periodic Markov model, is derived (Borodovsky and McIninch 1993). Similarly, the genomic segments labeled as ‘non-coding’ are used to estimate parameters for the non-coding model, structured as a 2^nd^ order uniform Markov chain.

#### The second iteration

##### Prediction step

The Viterbi algorithm is executed to produce the updated genome segmentation. Note that at this iteration we use the newly derived *typical* model along with all the *atypical* models.

##### Estimation step

The re-predicted genome segmentation is used to re-estimate parameters of the *typical* model. At this time, having information on initially predicted gene starts, the algorithm selects sequences situated around the gene starts and derives models of the patterns encoding transcription and/or translation regulation (Fig. S1).

#### Building the models of sequences around gene starts

In general, the sites near the first genes in operons (FGIOs) regulate transcription and translation (promoters and RBS sites) while the sites located near the interior genes in operons (IGIOs) regulate translation (RBS sites only). Given a predicted genome segmentation into protein-coding and non-coding regions, we identify FGIOs by the following rule. A gene is an FGIO if its upstream gene neighbor is located in the same strand at a distance larger than 25 nt or in the complementary strand (at any distance). The set of FGIOs is updated at each iteration. The rational for choosing the 25 nt threshold was as follows. Intuitively, larger values offer a more conservative selection of operons and FGIO, since distances between operons tend to be larger than distances between genes in operons. The operons annotated in the *E. coli* genome (Gama-Castro et al. 2016) were used to assess the effect of the threshold value on the accuracy of operon delineation. The 25 nt cut-off identified 98% of the annotated FGIOs correctly, while making ∼8% false positive predictions. Further, although 40 nt threshold could offer a slightly better balance between true and false operon predictions (Fig. S2), the experiments showed that the 40 nt threshold produced an equal (or even slightly worse) gene and gene-start prediction accuracy.

The two generic components of a regulatory site model, the nucleotide frequency matrix and the spacer length distribution, are then derived by running the modified Gibbs sampler algorithm GibbsL (described below). In some situations, it is useful to assess the “validity” of the motif by examining the preferred localizations of the motif instances. To do this, we define the *localization distance* as the mode of the spacer length distribution, i.e. the most frequent distance between the motif and the gene start. If the frequency at the mode is larger than a threshold value Q%, then the motif is considered to be *localized*.

The full details of the derivations of the models (for groups A-D, and X) illustrated by the ‘logos’ of positional frequency patterns of the motifs and the associated spacer length distributions (Figs. S3-S7) are given in the Supplement Materials

##### The third iteration and on

From now on, the *type* of the model of the gene start (and, therefore, the group to which the genome is supposed to “belong” to) does not change. However, the model parameters are updated in iterations with respect to the type-specific rules. At each prediction step, the dual motif models of the RBS and promoter box models (defined for both groups C and D) compete in the Viterbi algorithm computations. Similarly, the RBS and extended upstream signature models compete when the group X genome is processed. The *typical* gene model is updated at each iteration (Fig. 2) using the latest version of the genome segmentation.

GeneMarkS-2 continues the prediction/estimation iterations until the convergence condition is fulfilled (a 99% identity in gene starts between consecutive iterations). Alternatively, the algorithm stops if the maximum number of iterations is reached (10 by default). All the sequence segments labeled as coding regions in the final iteration are reported as predicted genes.

### A motif finder that accounts for the signal localization pattern

The MCMC motif finder Gibbs3 (Thompson, Rouchka, and Lawrence 2003) was designed to learn a probabilistic model of an *a priori* unknown motif present in a set of sequences. Gibbs3 was used for the RBS model delineation in GeneMarkS with reasonable accuracy (Besemer, Lomsadze, and Borodovsky 2001).

It was observed that sequences separating regulatory motifs from gene starts (spacers) have some preferred (more frequent) lengths, presumably facilitating molecular interactions involved in translation initiation. However, the Gibbs3 algorithm treats motif instances with spacer lengths that are too long or too short in the same way as the motif instances having a more optimal spacer length.

Therefore, we implemented a modified Gibbs sampler type algorithm, called GibbsL (see Supplemental Materials). We explicitly included the spacer length into the objective function in order to penalize motifs that appear at distances that vary significantly from the optimal locations. At a given iteration of GeneMarkS-2, GibbsL runs a fixed number, N, of its own iterations to derive a full model for a regulatory site (default N=60). Furthermore, the instances of GibbsL runs are repeated M times to mitigate the effect of random initializations (default M=30); then, the result with the highest objective function value is selected.

#### Test set-genes supported by proteomic studies

Mass-spectrometry-determined peptides were obtained in studies of a number of prokaryotic species at the Pacific Northwest National Laboratory (Venter, Smith, and Payne 2011). From all the available genomes, we selected 54, each with more than 250 proteomics validated ORFs (supported by at least two matching peptides). The peptide-supported ORFs (psORFs) annotated in the 54 genomes (Table S1) were used in the assessment of false negative and false positive rates of gene prediction.

#### Test set-COG annotated genes

We used 145 genomes (115 bacteria and 30 archaea covering 22 bacterial and archaeal phyla, listed in Table S2) suggested by our colleagues at DOE Joint Genome Institute (N. Kyrpides, personal communication). The genomes varied in size, type of genetic code, and GC content. Among the genes annotated in these genomes we selected genes with the COG characterization, therefore, the orthologous relationships of their protein products with proteins from other species were earlier established within the Clusters of Orthologous Groups (Tatusov, Koonin, and Lipman 1997; Tatusov et al. 2003; Galperin et al. 2015). The COG annotation serves as a robust evidence of the functional role of a gene, thus this gene set is unlikely to include random ORFs. Since 36 out of 145 genomes belonged to the set of 54 genomes with ‘proteomics’ confirmed genes, we removed the redundancy in the actual tests (see below).

#### Test sets with simulated genome specific non-coding regions

*Annotated* intergenic regions of each of 145 genomes (Table S2) we used to build models of non-coding sequence (zero-order Markov chains). Notably, such a simple type of model of the non-coding region was not used in any of the four gene finders. We employed these models to generate species specific random non-coding sequences and used them as follows. Firstly, in each genome out of 145 mentioned above (excluding *Mycobacterium leprae* known for large number of pseudogenes) we replaced intergenic regions by simulated non-coding sequences of the same length as in the original genome (Set 1). Secondly, we employed the models to generate 145 sequences of length 100,000 nt (Set 2). Thirdly, we added 100,000 nt of simulated non-coding sequence to the 3’ end of each genome (Set 3).

#### Test sets of genes with experimentally verified starts

N-terminal protein sequencing is a standard technique to validate sites of translation initiation (protein N-terminals and gene starts). Relatively large sets of genes with validated starts were known for the bacteria *Synechocystis sp.* (Sazuka, Yamaguchi, and Ohara 1999), *E. coli* (Rudd 2000; Zhou and Rudd 2013), *M. tuberculosis* (Lew et al. 2011), and *D. deserti* (de Groot et al. 2014) and the archaea *A. pernix* (Yamazaki et al. 2006), and *H. salinarum, N. pharaonis* (Aivaliotis et al. 2007).

#### Set of representative prokaryotic genomes

The prokaryotic genome collection of NCBI includes a description of 5,007 species as ‘representatives’ of the whole database of more than 100,000 genomes (Tatusova et al. 2014). These include 238 archaeal and 4,769 bacterial species to cover all the genera while leaving out the majority of species of the respective genera along with most of their strains.

Detailed descriptions of the data sets used in this paper are available either in the Supplemental Materials or at the site http://topaz.gatech.edu/GeneMark/GMS2/

## Results

### Error rates in prediction of protein-coding genes (all but the 5’ end)

Gene predictions made by GeneMarkS, Glimmer3, Prodigal, and GeneMarkS-2, run with default settings, were compared with (i) annotation of the MS or ‘proteomics’ validated genes and (ii) the COGs validated genes. In the 54 genomes (Table S2), there were ∼89,500 proteomics supported genes (psORFs, Table 2); in the 145 genomes (Table S3), there were ∼341,486 genes in total, 287,237 of which did not overlap with the proteomics validated genes (Table 2, 3).

In the set of 54 genomes, we observed that GeneMarkS-2 missed 181 psORFs out of 89,466, the least number of false negative errors made by the tested tools (Table 2). At the same time, GeneMarkS-2 made the least number of false positive predictions, 114. A predicted gene was judged as false if more than 30% of its length overlapped with a psORF located in one of five other frames.

**Table 2.** Statistics of false negative (panel A) and false positive gene predictions (panel B) observed in tests on 54 genomes containing proteomic validated genes and on 145 genomes with genes validated by orthologues in COGs.

| **A** |  |  |
| --- | --- | --- |
| Algorithm | Missed MS confirmed genes (from 89,466) | Missed COG genes (not MS) (from 287,237) |
| GeneMarkS | 376 | 1,467 |
| Glimmer3 | 496 | 1,990 |
| Prodigal | 217 | 1,389 |
| GeneMarkS-2 | **181** | **1,147** |

| **B** |  |  |
| --- | --- | --- |
| Algorithm | False predictions overlapping MS confirmed genes | False predictions overlapping COG genes (not MS) |
| GeneMarkS | 352 | 2,046 |
| Glimmer3 | 921 | 6,435 |
| Prodigal | 211 | 1,339 |
| GeneMarkS-2 | **114** | **932** |

**Table 3.** Panel **A**: Counts of genes missed by a particular tool (*false negatives*) among 341,486 COG genes annotated in 145 genomes. The counts are given in five length bins. Panel **B**: Counts of *false positive* predictions made in 144 simulated genomic sequences made from 144 original genomes where annotated intergenic regions were replaced by artificial non-coding sequence (see text). The numbers of false predictions were sorted by length in the same way as in Panel A.

| **A** | Bins (nt): | < 150 | 150-300 | 300-600 | 600-900 | > 900 | Total |
| --- | --- | --- | --- | --- | --- | --- | --- |
| Algorithm | COG genes | 362 | 13,985 | 65,948 | 83,745 | 177,446 | 341,486 |
|  |  | Missed annotated genes (FN) |  |  |  |  |  |
| GeneMarkS |  | 136 | **494** | 434 | 192 | 296 | 1,552 |
| Glimmer3 |  | **66** | 678 | 1,170 | 341 | 323 | 2,578 |
| Prodigal |  | 161 | 639 | 417 | 92 | 78 | 1,387 |
| GeneMarkS-2 |  | 132 | 596 | **370** | **76** | **69** | **1,243** |

**Table 3.** Panel A: Counts of genes missed by a particular tool (*false negatives*) among 341,486 COG genes annotated in 145 genomes.
| **B** | Bins (nt): | < 150 | 150-300 | 300-600 | 600-900 | > 900 | Total |
| --- | --- | --- | --- | --- | --- | --- | --- |
| Algorithm |  | False positives (FP) in simulated sequence |  |  |  |  |  |
| GeneMarkS |  | 3,366 | 5,113 | 1,230 | 177 | 94 | 9,980 |
| Glimmer3 |  | 17,446 | 5,044 | 1,299 | 228 | 136 | 24,153 |
| Prodigal |  | 4,525 | 5,321 | 1,453 | 419 | 135 | 11,853 |
| GeneMarkS-2 |  | **792** | **1,541** | **601** | **137** | **77** | **3,148** |

The comparison of the predictions with the COG validated genes demonstrated higher accuracy of GeneMarkS-2 as well. The new tool missed the lowest number of COG genes, 1,147, followed by Prodigal with 1,389. Notably, the rate of missed COG genes by any single gene finder was less than 1% (Table 2). Counting false positives (identified also as ones with prohibitively long overlaps with verified genes) has shown that GeneMarkS-2 made 932 false predictions, a significantly smaller number than the ones made by the other gene finders (Table 2). Note that the COG validated genes identical to the ‘proteomics’ supported genes were excluded from the second test.

We specifically looked into the distributon of the two types of errors with respect to the gene length (Table 3). Among all COG annotated genes Glimmer3 missed the least number of short genes (in 90-150 nt range) in comparison with the other tools (Table 3A). In the next bin, 150-300nt, GeneMarkS did show the best result. We observed that GeneMarkS-2 missed the least numbers of COG genes with length > 300nt and made the least total count (Table 3A). Its performance was the least dependent on genome GC content (Fig. S8).

Since it turned out that the false positives identified by their too long overlaps with validated genes (confirmed by proteomics or by COG annotation) occurred in rather small numbers (Table 2), we attempted to offer more substantial statistics by adding tests on sets of synthetic sequences.

### False positive predictions in synthetic sequences

We estimated the false positive rates on the three sets described above. First, we ran the four gene finders on Set1, the 144 constructs where the intergenic sequences were replaced by the same length synthetic non-coding sequences while the annotated genes remained in place. The predicted genes with 3’ end not matching annotation were considered as false positives (Table 3B). Notably, the number of false negatives in these experiments was observed to be of the same order as in the runs of the gene finders on the original genomes, i.e. ∼1% of the number of annotated genes. Second, we ran the four gene finding tools trained on 145 complete genomes on Set 2, the 145 sequences of length 100,000 nt, the experiment repeated 10 times (Table S5). The first and the second types of experiments had the advantage of keeping the trained parameters consistent with the non-perturbed features of the genome. However, the gene finders were not adapted to the simulated non-coding sequences. Therefore, we ran the gene finders in full cycle, training and prediction, on Set 3 where predictions made in the 100,000nt extended artificial portion of each genome were counted as false positives (Table S6).

The results of all the three types of tests described above were favorable for GeneMarkS-2. Further analysis demonstrated that reduction of false positives GeneMarkS-2 in comparison with GeneMarkS was mainly due to the improvement of the parameterization of the atypical models. Glimmer3 has made frequent false positives predictions of the short length (<150nt). This outcome is, arguably, the cost of the higher than other gene finders sensitivity in this length range (Table 2A).

Prodigal generated false positive predictions of rather long length as the algorithm settings give high weights to longer ORFs. Subsequently it leads to increase of false positives in genomic sequences with high GC content (Fig. S8A) where longer ORFs appear more frequently than in low GC genomes.

All over, in all the five approaches of the assessment of false positive rates (two in natural sequences and three in artificial ones) we saw that GeneMarkS-2 demonstrated the best performance. Notably, it yielded the lowest numbers of false positives in all the length intervals.

### Accuracy of gene start prediction

As it was stated above, we collected data on genes with experimentally validated translation initiation starts. Currently available collections were limited to the species of *A. pernix, D. deserti, E. coli, H. salinarum, M. tuberculosis, N. pharaonis*, and *Synechocystis sp.* To assess the accuracy of gene start prediction we compared positions of predicted and annotated starts of these verified genes.

The observed error rate of GeneMarkS-2 was 4.4%, followed by Prodigal at 6.1%, GeneMarkS at 10.2% and finally, Glimmer3 at 13.2%. Therefore, GeneMarkS-2 made the largest number of correct predictions among the four gene finders. (Table 4).

**Table 4.**
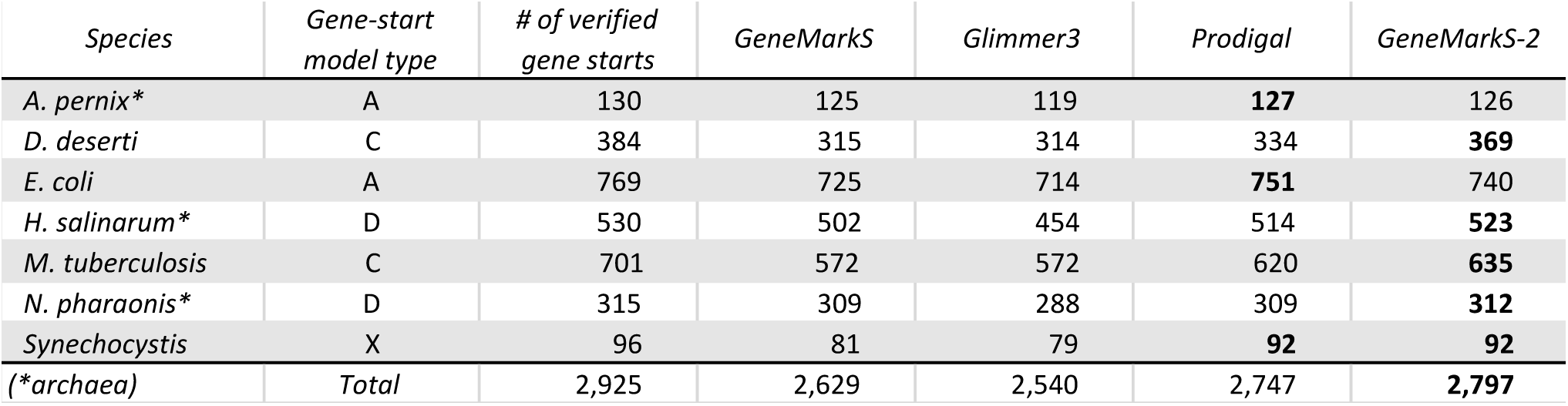
Numbers of correctly predicted gene starts verified by N-terminal protein sequencing.

In comparison with GeneMarkS, the improved start prediction accuracy of GeneMarkS-2 was due to a more flexible modeling of the regulatory signals near gene starts. For instance, in the group C genome of *M. tuberculosis*, GeneMarkS did not find a sufficiently strong RBS motif (Fig. 3A). GeneMarkS-2, on the other hand, predicted that 40% of operons are likely to be transcribed in the leaderless fashion, with the promoter Pribnow box located at a 6-8 nt distance from the gene starts (Fig. 3B). In the remaining ∼60% of operons, predicted RBS sites were separated from the gene starts by the same 6-8 nt distance (Fig. 3D). Therefore, the mixture of patterns, promoters and RBS sites, localized on the same distance from gene start was the reason for GeneMarkS to fail to converge to an informative motif model.

**Figure 3.**
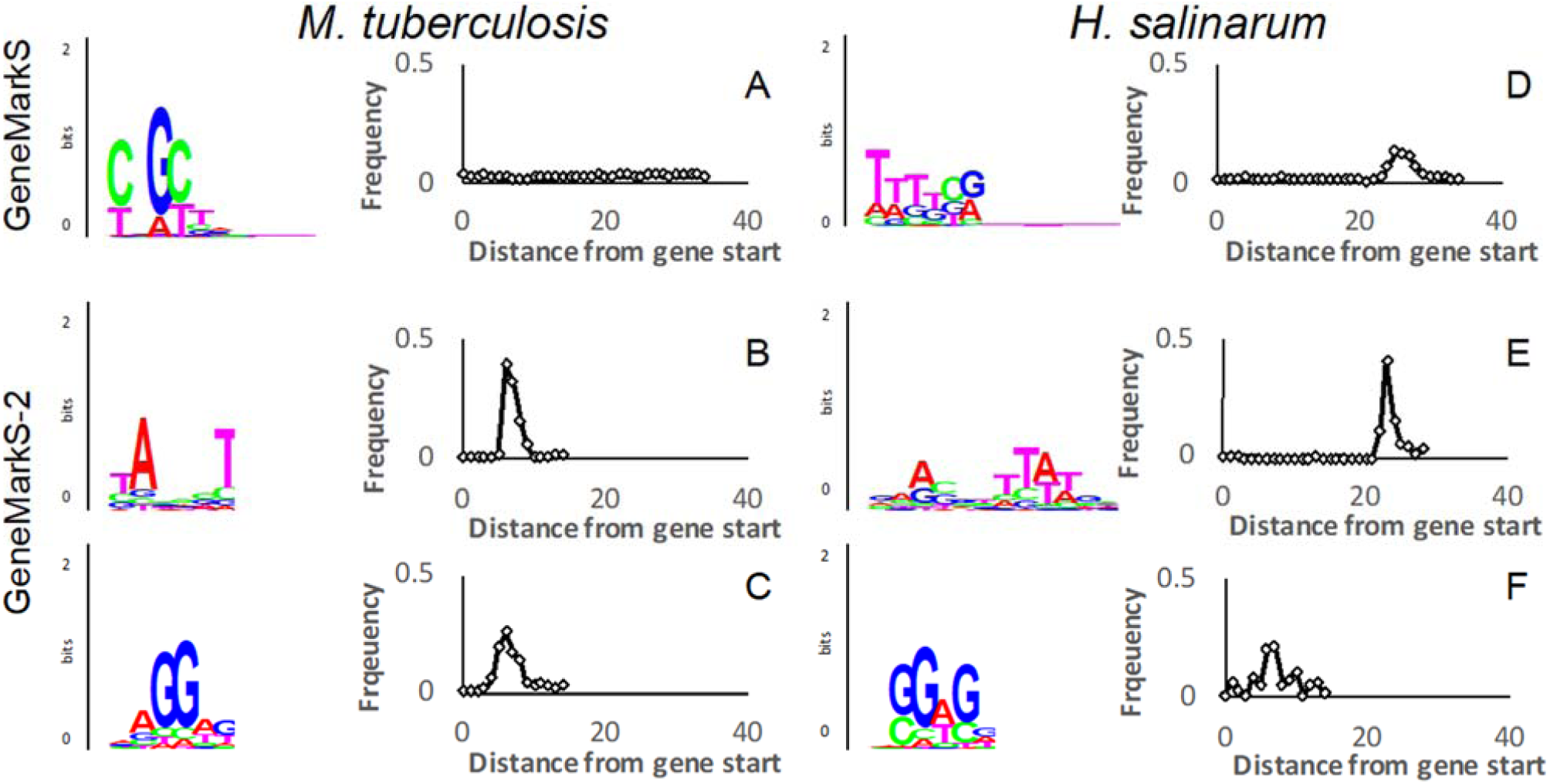
Motif logos and spacer length distributions for genomes of *M. tuberculosis* (**group C**) and *H. salinarum* (**group D**). In *M. tuberculosis*, the ‘mixed’ motif found by GeneMarkS has no preferred localization (panel A) in the upstream regions of the first genes in operons (FGIOs). To the contrary, the motif found by GeneMarkS-2 has a clear localization at -10 distance from gene starts, the distance typical for bacterial TATA box and leaderless transcription (B). In upstream regions of internal genes in operons (IGIO) GeneMarkS-2 built the RBS model and the spacer length distribution (C). For *H. salinarum* we comparison of GeneMarkS-2 outcomes (E and F) with results shown by GeneMarkS (B) shows similar improvements.

In another example, for majority of the first genes in operons in the group D archaea *H. salinarum* GeneMarkS-2 identified the promoter Pribnow box located at 22-24 nt from the gene starts (Fig. 3E), at the distance characteristic for leaderless transcription in archaeal genomes. For the remaining FGIOs as well as for IGIOs GeneMarkS-2 identified the RBS sites at 6-8 nt distance upstream to gene starts (Fig. 3F). GeneMarkS, that assumes a single type of motifs for all FGIOs, could only derive a Pribnow box like site motif with a weaker localization (Fig. 3D).

On a general note, the performance of GibbsL in detecting RBS motifs with the Shine-Dalgarno consensus in genomes with low and mid GC content was observed to be similar to the performance of Gibbs3 (though GibbsL tends to have higher localization peaks). However, in genomes with high GC content, GibbsL derived motifs with a higher information content and a more compact localization (Fig. S9, S10, S11). We also observed that GibbsL was more robust than Gibbs3 upon increasing the length of sequences (i.e. the selected gene upstream regions) supposed to contain common motifs (Fig. S9, S10).

Finally, we have observed that the selection of the width of the motif in GibbsL in the range from 5nt to 10nt, did not show a significant influence on gene start prediction (Table S3). Indeed, the motifs with larger widths, i.e. 10nt, show a minor change in comparison with the motifs with short width, e.g. 5nt, as shown in Fig S12.

### Analysis of ∼5,000 prokaryotic genomes with GeneMarkS-2

Upon generating gene predictions for ∼5,000 *representative* NCBI genomes (see above) GeneMarkS-2 made an assignment of each genome into one of the five groups/categories, A-D and X. We presented the species assigned to each category in Tables S4A, S4B,…S4X with names listed along the branches of five trees that follow the taxonomic order. Not surprisingly, the species from the same clades tend to belong to the same category. The distribution of the five categories at the top three taxonomical levels is shown in Fig. 4.

**Figure 4:**
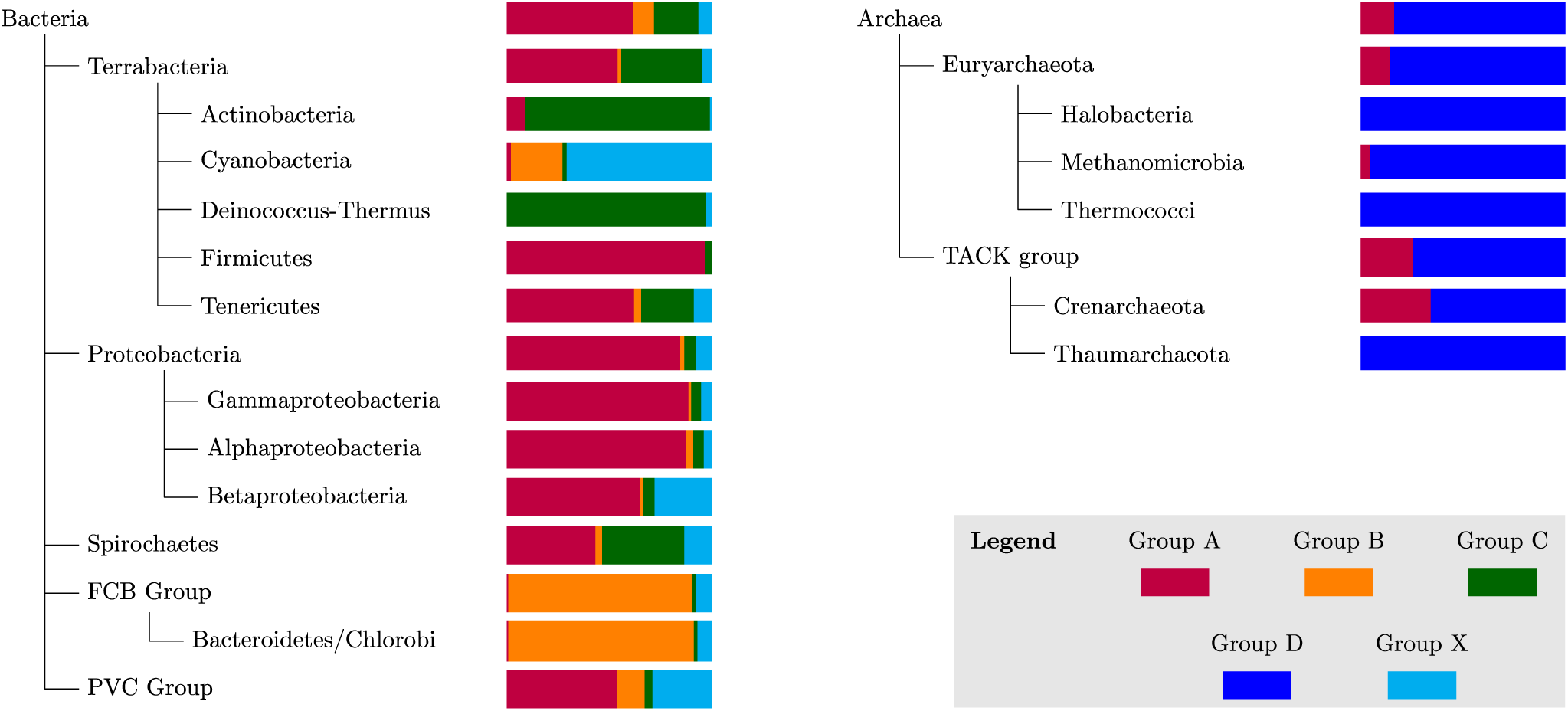
The distribution of groups A-D and X among ∼5000 representative genomes as determined by GeneMarkS-2. This diagram shows the top three levels of the taxonomy trees of both archaea and bacteria.

The largest, group A, includes 2,935 bacteria and 39 archaea (Table S4A). In the species of category A, gene expression occurs predominantly via mRNAs which 5’ UTRs carry detectable RBS motifs with the Shine-Dalgarno consensus. Among the category A bacteria, 61% were Gram-negative and 39% were Gram-positive. Still, these Gram-positive species in group A made more than half, 57%, of Gram-positive bacteria in the set of ∼5000 species. Furthermore, it is only Gram-positive *Actinobacteria* that rarely belong to group A (78 out of 859, 9.1%) and mostly appear in group C; 96% of the all other Gram-positive bacteria belong to group A.

The group/category B assignments were made for 495 bacteria and no archaea. The characteristic feature of this category is *non-SD type RBS* motif. In genomes of group B we observed the presence of the same type motif in the upstream sequences of all the genes, both initial and internal genes in operons, however, the motif did not have the Shine-Dalgarno consensus. Since this motif is present at a short distance from the translation start in the leadered mRNAs, it cannot be a promoter. Species of group B are frequent in the FCB group of bacteria (409 out of 455, 89.9%), but rare in *Terrabacteria* (1.7%) and *Proteobacteria* (2.0%).

Group C (1028 out of 4769 bacteria) consists of bacterial species predicted to have frequent presence of leaderless mRNAs (Table S4C). Species of group C are frequent in *Actinobacteria* (773 from 859, 90.0%) and *Deinococcus-Thermus* (37 out of 38, 97.4%), but rare in *Proteobacteria* (104 out of 1854, 5.6%) and *Firmicutes* (36 out of 1064, 3.4%). A particularly high frequency of category C species was observed in *Streptomycetales* (129 out of 129, 100%) and in *Corynebacteriales* (197 out of 202, 97.5%) including *Mycobacteriaceae* (56 out of 57, 98.2%).

Next is group D that includes archaeal species with prevalence of leaderless mRNAs. A promoter box like motif was derived in genomes of this category for leaderless FGIO, while for the remaining genes an RBS motif was determined. From the 238 archaeal genomes in our dataset, 199 were assigned to group D. In particular, some taxa had most (or all) of their members belonging to this group, such as *Halobacteria* (74 out of 74 species, 100%), *Methanomicrobia* (40 out of 42, 95%), *Thermococci* (21 out of 21, 100%), *Thermoplasmata* (11 out of 11, 100%), *Archeoglobi* (7 from 7, 100%), *Thaumarchaeota* (11 from 11, 100%) and *Crenarchaeota* (23 out of 35, 65%) (see Table S5D). Leaderless transcription is the characteristic feature of group D genomes. We have inferred, however, that group A is home for a significant fraction of the taxon *Crenarchaea*, where *P. aerophilum* belongs. Thus, many members of *Crenarchaea* should have a low percentage of leaderless transcripts.

Finally, 311 bacterial species did not fit any of the above four groups (A through D) and were included into group X (Table S5X) characterized by the (seeming) absence of pronounced regulatory signals upstream to most genes. Still, the weakness of the regulatory signal has its own commonality. Species of this group are relatively frequent in *Cyanobacteria* (90 out of 127, 70.9%) and in *Burkholderiales* (63 out of 166, 37.9%).

The summary list of the distribution of the ∼5,000 species among groups A-D, and X is given in Table 5.

**Table 5.** Distribution of archaeal and bacterial genomes among groups A-D, and X.

|  | Archaeal Genomes |  | Bacterial Genomes |  | Total |
| --- | --- | --- | --- | --- | --- |
|  | Number | % | Number | % | Number |
| Group A | 39 | 16.4 | 2,935 | 61.5 | 2,974 |
| Group B | 0 | 0.0 | 495 | 10.4 | 495 |
| Group C | NA | NA | 1,028 | 21.6 | 1,028 |
| Group D | 199 | 83.6 | NA | NA | 199 |
| Group X | 0 | 0.0 | 311 | 6.5 | 311 |
| Total | 238 | 100 | 4,769 | 100 | 5,007 |

## Discussion

### Gene finding accuracy evaluation

We demonstrated in several tests that, on average, GeneMarkS-2 is a more accurate tool than the current frequently used gene finders. Particularly, GeneMarkS-2 made fewer *false negative* and *false positive* errors in predicting genes validated by mass-spectrometry and COG annotation (Table 2). Also, the numbers of *false positive* predictions made by GeneMarkS-2 in simulated non-coding sequences were significantly smaller than the numbers observed for other tools (Table 3B).

The array of atypical models employed in GeneMarkS-2 improved the prediction of horizontally transferred (atypical) genes. In our observations, the deviation of GC composition of atypical genes from the genome average could be as large as 16% (e.g. the 798 nt long *E. coli* gene *b0546* characterized as DLP12 prophage, with GC content 36% compared to the 52% GC content of the bulk of *E. coli* genes). The GC content of atypical genes is frequently lower than the GC content of ‘typical’ ones (Fig. S13). Also, the ‘atypical’ genes with large GC content deviations are expected to appear more frequently in high GC genomes given the larger space for downward variation. All in all, atypical genes may constitute a significant fraction of the whole gene complement (e.g. about 15% of genes in the *E. coli* genome (Borodovsky et al. 1995)). In our analysis of the ∼5,000 genomes, we found that the distribution of the fraction of predicted atypical genes in prokaryotic genomes are rather similar between archaea and bacteria, with an average of about 8-9% (Fig. 5).

**Figure 5.**
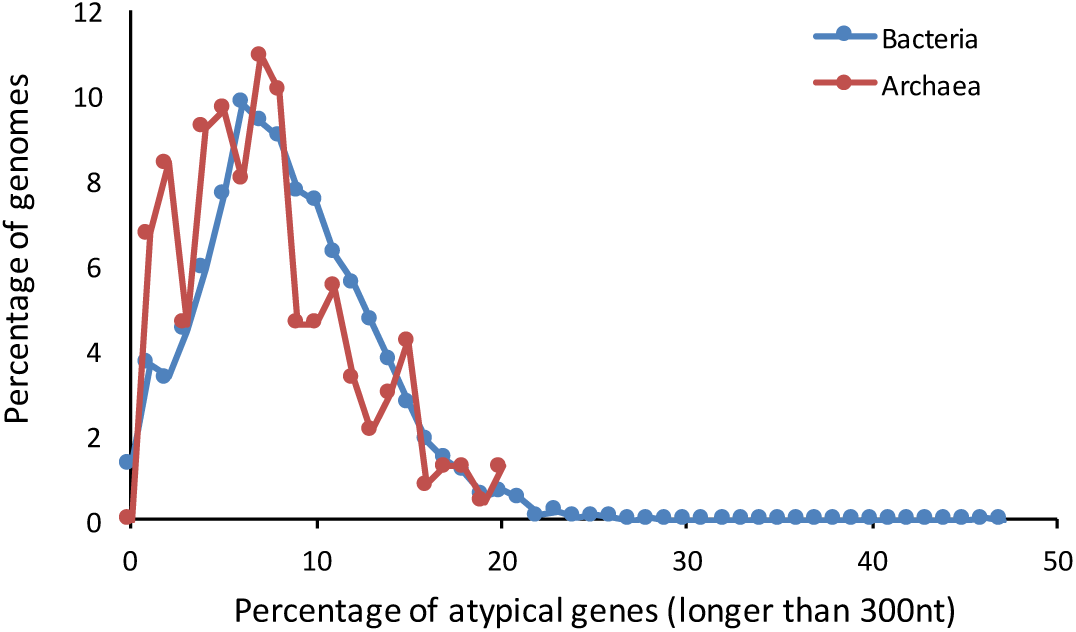
Distributions of the percentage of predicted atypical genes in archaeal and bacterial genomes.

A comparison of the sets of COG annotated genes missed by the three gene finders (Fig. S14) shows that atypical genes, genes predicted by atypical models of GeneMarkS-2, constituted 30% of 780 (534+246) genes missed by Prodigal and 42% from 1605 (1359+246) genes missed by Glimmer3. Both Prodigal and Glimmer3 employ a single model of protein-coding regions. We argue that the more accurate prediction of atypical genes by GeneMarkS-2 makes a compelling argument in favor of the use of multiple models of protein-coding regions.

One of the features of GeneMarkS-2 is the ability to characterize atypical genes as bacterial or archaeal due to the division of the *atypical* models into distinct bacterial and archaeal types (Zhu, Lomsadze, and Borodovsky 2010). The insights into the possible origin of atypical genes (likely horizontally transferred) could be particularly useful for genomes of thermophilic bacteria and mesophilic archaea.

GeneMark-S2 made the least number of errors in gene start predictions (120 out of 2,927), with the second-best tool, Prodigal, coming in at 170 errors (Table 4). Notably, Prodigal performed better on *E. coli* set, however, this set is special as it was used in the supervised training of Prodigal’s gene-start prediction model (Hyatt et al. 2010). As expected, GeneMarkS-2 made more accurate predictions for genomes with non-canonical RBS sites (group C) and archaeal genomes with frequent leaderless transcription (group D). Interestingly, the experimental study of *D. deserti* identified 384 genes with verified translation starts, 262 of which had transcription starts annotated with dRNA-Seq (de Groot et al. 2014). It was experimentally shown that 167 out of the 262 genes had leaderless transcription. In this genome, GeneMarkS-2 correctly predicted 34 more starts than Prodigal. While Prodigal only detects RBS motifs for this genome, GeneMarkS-2 builds both a promoter and a RBS model.

### Characterization of the patterns around gene starts. Prediction of the extent of leaderless transcription

GeneMarkS-2, the first among gene finding tools, is able to infer the presence of leadered or leaderless transcripts from the types of the regulatory motifs predicted upstream to the gene starts.

Screening made by GeneMarkS-2 allows to conclude that leaderless transcription is a ubiquitous feature in prokaryotic genomes; also, this screening allows to determine the extent of leaderless transcription within genomes (Fig. 6). In large number of analyzed archaeal genomes we predicted leaderless transcription of 60% to 80% of the operons. Still, in some archaeal species this fraction was only 25-35%. This fraction is closer to what was observed in bacterial genomes assigned to group C (25% to 50% of operons).

**Figure 6.**
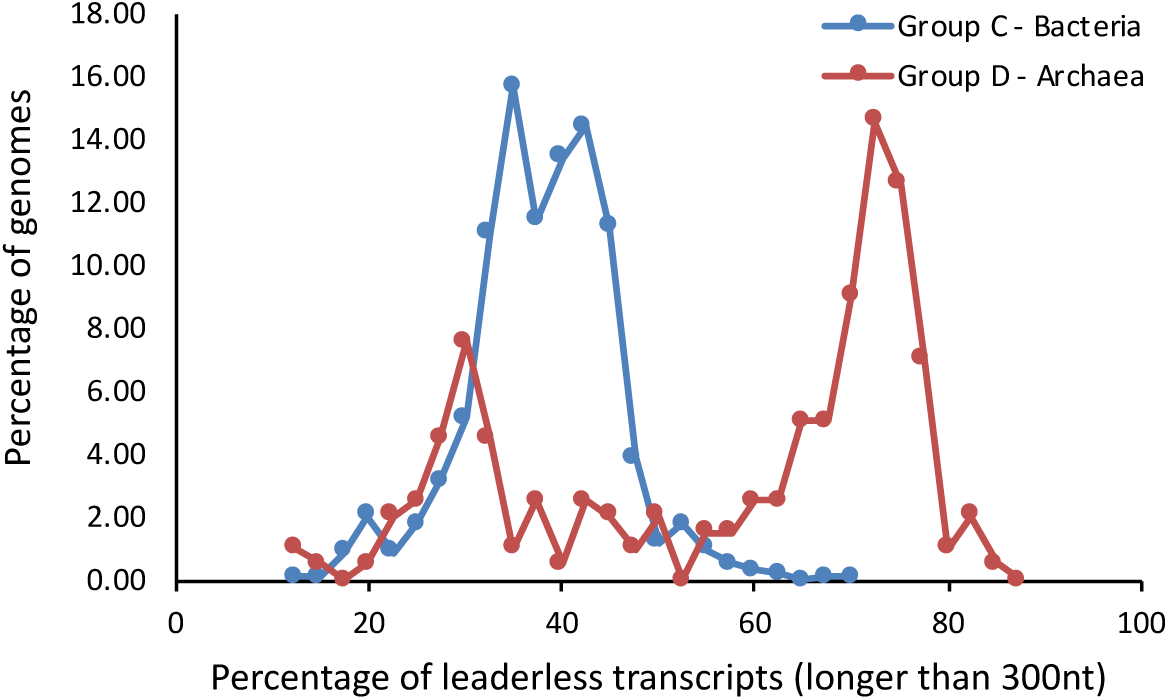
Distributions of the percentage of leaderless transcripts among all transcripts in bacterial group C and archaeal group D.

We have compared the predictions of the fractions of the leaderless transcripts with the observations made in the dRNA-Seq experiments with *Deinococcus deserti* (de Groot et al. 2014), *Haloferax volcanii* (Babski et al. 2016), *Sulfolobus solfataricus* (Wurtzel et al. 2010) and *M. tuberculosis* (Cortes et al. 2013).

For the sets of genes having identical predicted and experimentally determined 5’ and 3’ ends we found the following: in archaea, *D. deserti*: ∼62% predicted leaderless vs 62% observed leaderless (1,707 genes in total); *H. volcanii*: 86% vs 82% (1,406 genes); *S. solfataricus*: 78% vs 76% (859 genes); in bacteria, *M. tuberculosis*: 42% vs 34% (1310 genes). Thus, the predictions were in reasonably good agreement with the experiment.

Genomes with experimentally characterized small numbers of leaderless genes (Sharma et al. 2010; Thomason et al. 2015; Kroger et al. 2013; Wiegand et al. 2013; Dugar et al. 2013; Nicolas et al. 2012; Shao et al. 2014; Pfeifer-Sancar et al. 2013) (TSS experiments) were all classified as group A. Genomes with large proportion of leaderless transcripts were all classified as group B (Cortes et al. 2013; de Groot et al. 2014; Shell et al. 2015; Pfeifer-Sancar et al. 2013) or group D (Koide et al. 2009; Babski et al. 2016; Wurtzel et al. 2010; Jager et al. 2014; Toffano-Nioche et al. 2013).

Experiments on *Synechocystis sp* demonstrated the prevalence of leadered transcription (Mitschke et al. 2011). However, GeneMarkS-2 was able to detect an RBS motif (with the SD consensus) in less than 15.5% of its genes. Experiments have shown that mutating some “A” rich sequences situated 15-45 nt upstream to the gene starts (still within the long 5’ UTR sequence) led to changes in gene expression (Mutsuda and Sugiura 2006). That said, the mechanism used for the translation start recognition in the majority of *Synechocystis sp* genes is unknown.

It was observed that translation initiations of the three types, SD-RBS based, non-SD RBS based, and leaderless are present in *E. coli* (Barrick et al. 1994; Shean and Gottesman 1992; Resch et al. 1996). Further observations have shown that the distribution of the numbers of genes controlled by each of the three mechanisms could vary significantly between species (Gualerzi and Pon 2015). If a particular type of translation initiation appears rarely in a given species, the GeneMarkS-2 training procedure is not able to make models for the motifs around gene starts regulated by this type of mechanism due to the insufficient size of the training set.

### Regulatory motifs in the species of group B

GeneMarkS-2 currently assigns 495 out of 4,769 bacteria (and none out of 238 archaea) to group B. Here we take as an example the genome of *Bacteroides ovatus*. Interestingly, while its 16S rRNA features the *bona fide* anti-Shine-Dalgarno pattern, the SD-matching sequences appear upstream to the start sites in only ∼3% of genes. Moreover, the A-rich sequences were observed in the upstream regions of the majority of *B. ovatus* genes. It was shown that mutating these A-rich regions reduces gene expression levels. Thus, the A-rich sequences are important for the translation start recognition (Wegmann, Horn, and Carding 2013). GeneMarkS-2 identified the A-rich non-SD type motif with consistent localization at ∼9 nt from TIS (Fig. 7).

**Figure 7.**
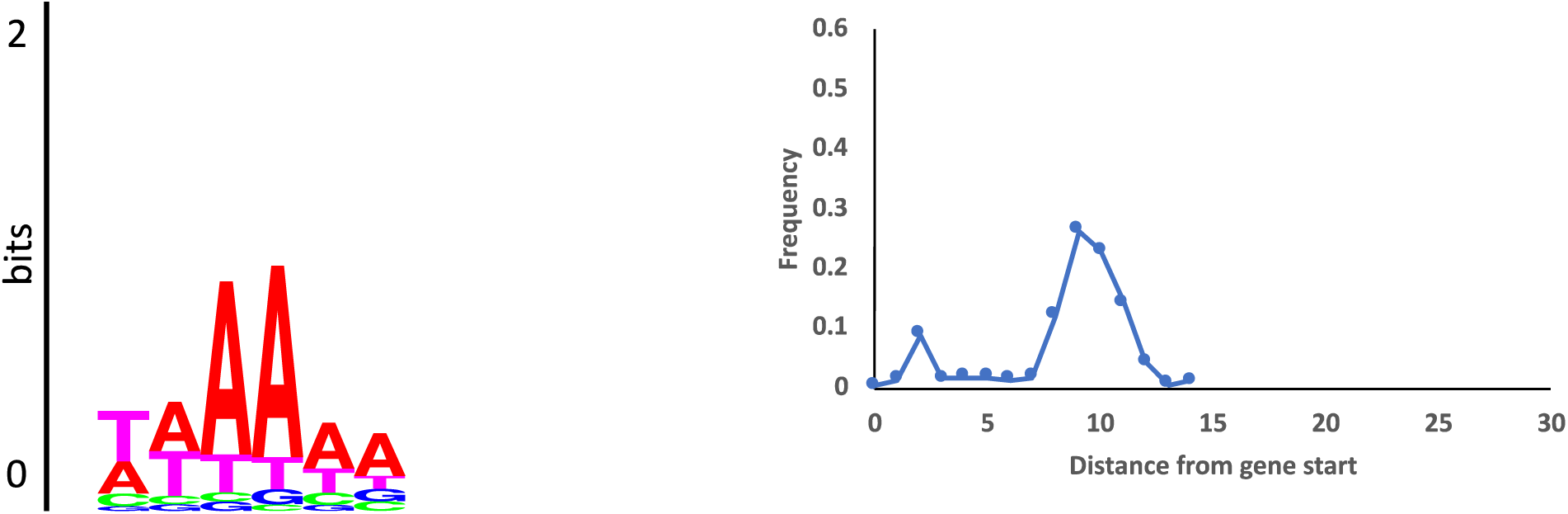
The motif logo and the spacer length distribution of *Bacteroides ovatus*, a group B genome.

Note that GeneMarkS-2 assigned 90% of *Bacteroidetes/Chlorobi* genomes to group B (408 of 450). While not much is known about the non-SD mechanisms in bacteria, the clustering of the species assigned to group B within specific parts of the taxonomy tree lends additional credibility to the results (Table S5, D).

Particularly, in *Bacteroides* (including *B. ovatus*) 21 out of the 23 genomes in this study were assigned to group B. In these 21 genomes, GeneMarkS-2 found motifs similar in conservation pattern and localization distribution to those revealed in *B. ovatus*.

Also, 30 out of the 30 *Flavobacterium* genomes (a genus from *Bacteroidetes/Chlorobi*) were assigned to group B. The 6 nt motifs, similar in the conservation pattern to the one from *B. ovatus*, were situated at the same distance from translation start sites (Fig. S15 for *Flavobacterium frigidarium*).

While genomes of the species from these genera are closely related, there are some differences in the derived motifs. In particular, *Bacteroides* tends to have a few strong A nucleotides next to the ‘core’ motif and close to translation start, hence the peak in the spacer length distribution at ∼3nt distance from the starts (Fig. S16A). In *Flavobacterium* the ‘core’ motif with consensus TAAAAA is better pronounced and its position better coincides with the motif 3’ end, hence the spacer length distribution has a peak at ∼7nt distance (Fig. S16B). The consistency of this observation was tested for all 21 *Bacteroides* and 30 *Flavobacteria*. The 6 nt core motif was not easy to detect in *Prevotella* (a close relative of *Bacteroides*) when the motif width was set to 6 nt. However, setting the motif width to 15 nt yielded a similar motif. Interestingly, unlike *B. ovatus*, other group B species may have 16S rRNA with a mutated or truncated tail (Lim, Furuta, and Kobayashi 2012).

In a recent publication (Nakagawa, Niimura, and Gojobori 2017), leaderless and non-SD initiation were included into the same class. Here, we made the distinction between the leaderless transcripts and, thus, the absence of RBS upstream to the first genes in operons (groups C and D) and the leadered transcripts with non-SD RBS motifs in 5’UTR (group B). GeneMarkS-2 is able to identify species that use non-SD RBS along with the detection of the motifs per se.

The remaining difficulties in automatic genome annotation are concerned with a minor fraction of genes in any given genome. We observed that just a small number genes may still escape detection by *ab initio* tools, e.g. genes significantly biased in higher order oligonucleotide composition or genes containing frameshifts. When an orthologue of such a gene is present in the database, the frameshift identification can be done rather easily. The database search and alignment helps even for functional frameshifted genes that are conserved in evolution, such as in the *prfB* gene encoding a translation initiation factor (Craigen and Caskey 1986) or genes regulating some mobile elements (Sharma et al. 2011). Furthermore, pseudogenes, especially expressed pseudogenes, could mislead gene finding tools into generating predictions that are difficult to classify as false positives.

An extension of GeneMarkS (known as GeneMarkS+), integrates external evidence, such as protein homology, into *ab initio* gene prediction. In the NCBI prokaryotic genome annotation pipeline GeneMarkS+ integrates several types of evidence into genome annotation (Tatusova et al. 2016). Similar extension is logical to make for GeneMarkS-2.

## Software and data availability

The following websites can be used to access the GeneMarkS-2 software: http://topaz.gatech.edu/GeneMark/genemarks2.cgi, and data used in the course of this research project: http://topaz.gatech.edu/GeneMark/GMS2/ Running time of GeneMarkS-2 is currently ∼ 3 minutes on a genome of a size of *E. coli*.

## Acknowledgements

We thank Susan M.E. Smith for useful comments on the manuscript. This work was supported by National Institutes of Health (NIH) [HG000783 to M.B.].

